# PhoPQ-mediated lipopolysaccharide modification regulates intrinsic resistance to tetracycline and glycylcycline antibiotics in *Escherichia coli*

**DOI:** 10.1101/2024.07.01.601565

**Authors:** Byoung Jun Choi, Umji Choi, Dae-Beom Ryu, Chang-Ro Lee

## Abstract

Tetracyclines and glycylcycline are among the last-resort antibiotics used to combat infections caused by multidrug-resistant Gram-negative pathogens. Despite the clinical importance of these antibiotics, their mechanisms of resistance remain unclear. In this study, we elucidated a novel mechanism of resistance to tetracycline and glycylcycline antibiotics via lipopolysaccharide (LPS) modification. Disruption of the *Escherichia coli* PhoPQ two-component system, which regulates the transcription of various genes involved in magnesium transport and LPS modification, leads to increased susceptibility to tetracycline, minocycline, doxycycline, and tigecycline. These phenotypes are caused by enhanced expression of phosphoethanolamine transferase EptB, which catalyzes the modification of the inner core sugar of LPS. PhoPQ-mediated regulation of EptB expression appears to affect the intracellular transportation of doxycycline. Disruption of EptB increases resistance to tetracycline and glycylcycline antibiotics, whereas the other two phosphoethanolamine transferases, EptA and EptC, that participate in the modification of other LPS residues, are not associated with resistance to tetracyclines and glycylcycline. Overall, our results demonstrated that PhoPQ-mediated modification of a specific residue of LPS by phosphoethanolamine transferase EptB regulates resistance to tetracycline and glycylcycline antibiotics.

**Importance:** Elucidating the resistance mechanisms of clinically important antibiotics helps in maintaining the clinical efficacy of antibiotics and in the prescription of adequate antibiotic therapy. Although tetracycline and glycylcycline antibiotics are clinically important in combating multidrug-resistant Gram-negative bacterial infections, their mechanisms of resistance are not fully understood. Our research demonstrates that the *Escherichia coli* two-component system PhoPQ regulates resistance to tetracycline and glycylcycline antibiotics by controlling the expression of phosphoethanolamine transferase EptB, which catalyzes the modification of the inner core residue of lipopolysaccharide (LPS). Therefore, our findings highlight a novel resistance mechanism to tetracycline and glycylcycline antibiotics and the physiological significance of LPS core modification in *E. coli*.

**One sentence summary:** Lipopolysaccharide modification-mediated tigecycline resistance

## INTRODUCTION

Bacteria encounter diverse environmental stresses such as exposure to antibiotics, which can cause serious growth arrest or cell lysis. In a single cell, bacteria must be equipped with all the stress response mechanisms necessary to overcome stress conditions. Therefore, bacteria have precise stress adaptation mechanisms such as the two-component system. The two-component system comprises of a sensor kinase and response regulator (1). Sensor kinase, an inner membrane protein with transmembrane helix domains, senses extracellular stress conditions. These conditions result in conformational changes in its structure and induces autophosphorylation at a specific residue of its cytoplasmic domain (1). Subsequently, the phosphate group in the cytoplasmic domain of the sensor kinase is transferred to a specific residue of the response regulator to induce conformational changes in the response regulator. Phosphorylated response regulators can induce or repress the transcription of various target genes (1).

Bacteria possesses several two-component systems that regulate a diverse range of stress responses. The PhoPQ system is an important two-component systems involved in adaptation to envelope stresses. In this system, the sensor kinase PhoQ phosphorylates the response regulator PhoP in response to various conditions, such as magnesium depletion (2), acidic stress (3), exposure to cationic antimicrobial peptides (4), and osmotic upshift (5). Phosphorylated PhoP regulates the transcription of several genes, such as the *mgtA* gene encoding a magnesium transporter that mediates the import of magnesium under magnesium starvation conditions (6, 7). The PhoPQ system also plays a pivotal role in the survival of intracellular bacterial pathogens such as *Salmonella enterica*, inside macrophage (8, 9). Although the PhoPQ system enables bacteria to survive under diverse environmental conditions, the regulatory mechanisms underlying PhoPQ-mediated antibiotic resistance remain unclear.

Tetracycline and its derivatives such as minocycline and doxycycline are used to treat infections caused by multidrug-resistant Gram-negative pathogens (10). Tigecycline is a unique antibiotic in the glycylcycline class derived from tetracycline, and is among the last-resort antibiotics for the treatment of severe infections caused by multidrug-resistant Gram-negative pathogens (11). Tigecycline was developed to evade several tetracycline resistance mechanisms, such as the ribosome protection mechanism (TetM) and efflux pump mechanism (TetA), via the addition of a t-butylglycylamido moiety at the 9^th^ position of the tetracycline ring D of minocycline, resulting in an increased affinity for ribosomes (11, 12). Owing to their clinical importance, the resistance mechanisms to tetracyclines and glycylcycline, such as efflux pumps, ribosome modifications, and the production of inactivation enzymes, have been investigated in several Gram-negative pathogens (10, 13–15). However, the impact of altered outer membrane permeability, such as lipopolysaccharide (LPS) modification, on the resistance to tetracycline and glycylcycline antibiotics is not fully understood.

In this study, we demonstrated that LPS core modification regulates resistance to tetracycline and glycylcycline antibiotics in *Escherichia coli*. The PhoPQ two-component system represses the expression of phosphoethanolamine transferase EptB which catalyzes the modification of the inner core sugar of LPS, which is necessary for resistance to tetracycline and glycylcycline antibiotics. Additionally, of the three phosphoethanolamine transferases (EptA, EptB, and EptC) involved in LPS modification, only EptB is associated with resistance to tetracyclines and glycylcycline. Overall, these results suggest that PhoPQ-mediated regulation of EptB expression affects resistance to tetracycline and glycylcycline antibiotics.

## RESULTS

### Inactivation of sensor kinase PhoQ induces increased susceptibility to minocycline and tigecycline

We constructed several mutants defective in sensor kinases associated with the envelope stress response to determine the effect of the two-component system on antibiotic resistance, and examined the MICs of various antibiotics with different modes of action in these mutant strains. Among the 11 mutant strains, six mutant strains did not exhibit any change in the MICs of the antibiotics tested (Fig. S1), whereas five mutant strains exhibited changes in the MICs for one or more antibiotics (Fig. 1). The MICs of β-lactams (blue bars), aminoglycosides (orange bars), and fosfomycin (yellow bar) were increased in the Δ*cpxA* mutant, compared to those in the wild-type strain (Fig. 1). A previous study reported that CpxA affects resistance to β-lactams via regulation of *ompF* and *slt* expression, aminoglycosides resistance via regulation of *acrD* expression, and fosfomycin resistance via regulation of *glpT* and *uhpT* expression (16). β-lactam resistance was also affected in the Δ*envZ* mutant (green bars) (Fig. 1). These results may be owing to altered expression levels of OmpF and OmpC porins, which strongly affect β-lactam resistance through the penetration of the outer membrane and the maintenance of membrane integrity (17). The MIC of fosfomycin was 4-fold higher in Δ*uhpB* mutant than in the wild-type strain (purple bar) (Fig. 1). Previous studies have revealed that UhpB affects fosfomycin resistance via regulation of the expression of a hexose phosphate transporter UhpT (18, 19). The MICs of several antibiotics were also changed in the Δ*phoQ* mutant (Fig. 1). Among them, we focused on the increased susceptibility to minocycline and tigecycline in the Δ*phoQ* mutant (red bars). Although minocycline and tigecycline are potent antibiotics in treating infections caused by multidrug-resistant Gram-negative pathogens (20, 21), their mechanisms of resistance remains poorly understood, which prompted us to elucidate the underlying mechanisms of these phenotypes.

**FIG 1.**
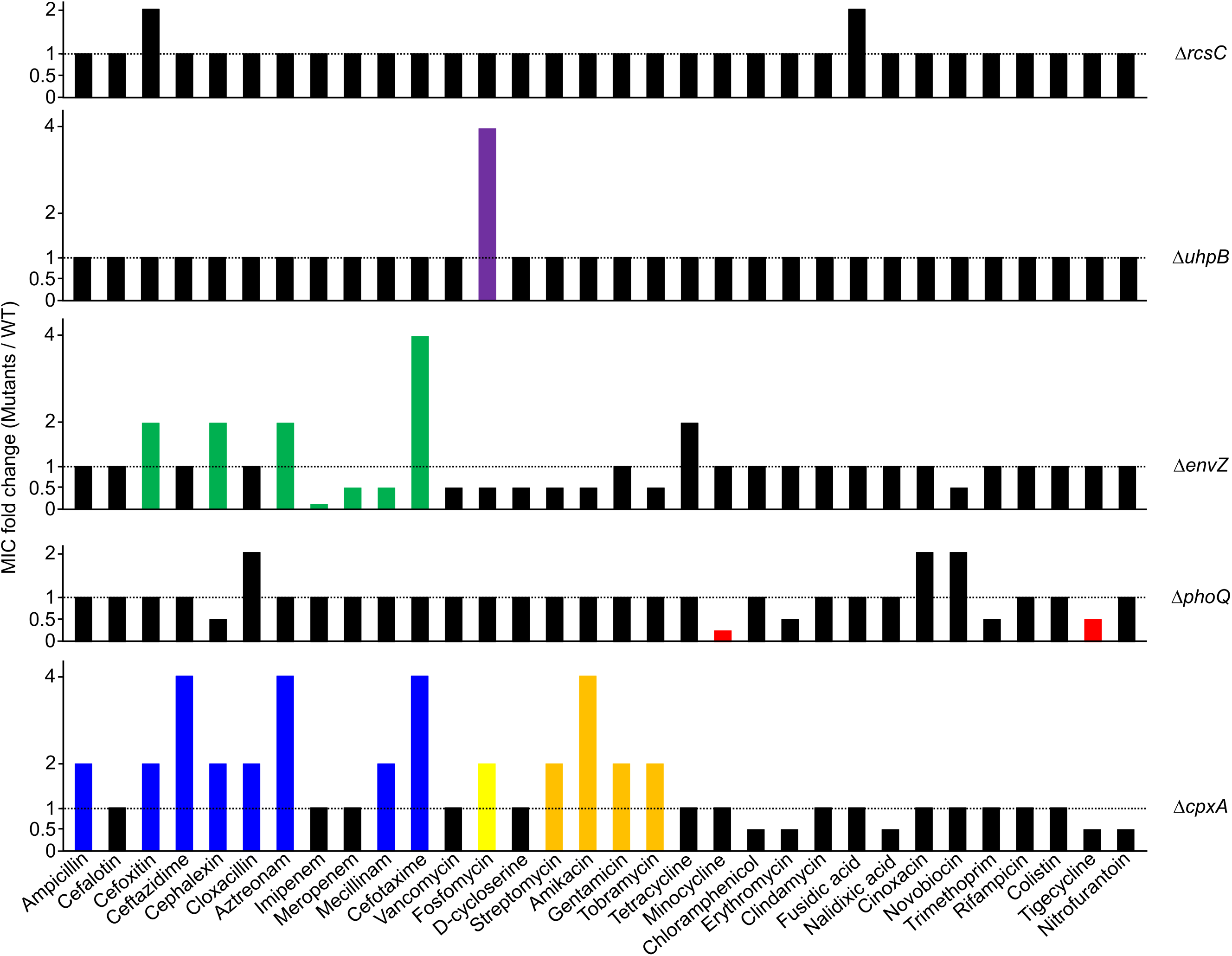
The inactivation of two-component system sensor kinases affects intrinsic antibiotic resistance. The MICs of various antibiotics were measured against the wild-type and indicated mutant strains in MH medium. The relative MIC values for the indicated mutant cells compared to those for the wild-type cells are presented. Colored bars indicate the MIC values of important antibiotics which increase or decrease in the indicated mutant cells.

### PhoPQ two-component system is required for resistance to tetracycline and glycylcycline antibiotics

The MIC of minocycline was 4-fold lower in Δ*phoQ* mutant than in the wild-type strain, but the other mutant strains did not exhibit any changes (Fig. 2A). Minocycline and doxycycline belong to the tetracycline class, whereas tigecycline is a member of the glycylcycline class, which is derived from tetracycline (12). Therefore, the structural differences among these antibiotics (tetracycline, minocycline, doxycycline, and tigecycline) were not significant (Fig. 2B). The MICs of doxycycline, tigecycline, and tetracycline in the Δ*phoQ* mutant were 4-fold, 2-fold, and 1.5-fold lower, respectively, than in the wild-type strain (Figs. 2C and S2). We examined the effect of PhoP on antibiotic resistance as the sensor kinase PhoQ acts along with the response regulator PhoP. The MICs of the four antibiotics in the Δ*phoP* mutant were identical to those in the Δ*phoQ* mutant (Fig. 2C). The antibiotic sensitivity of the Δ*phoP* mutant was complemented by the pACYC184 plasmid-based expression of the *phoP* gene (Fig. 2D). These results indicate that the PhoPQ two-component system is necessary for intrinsic resistance to tetracycline and glycylcycline antibiotics.

**FIG 2.**
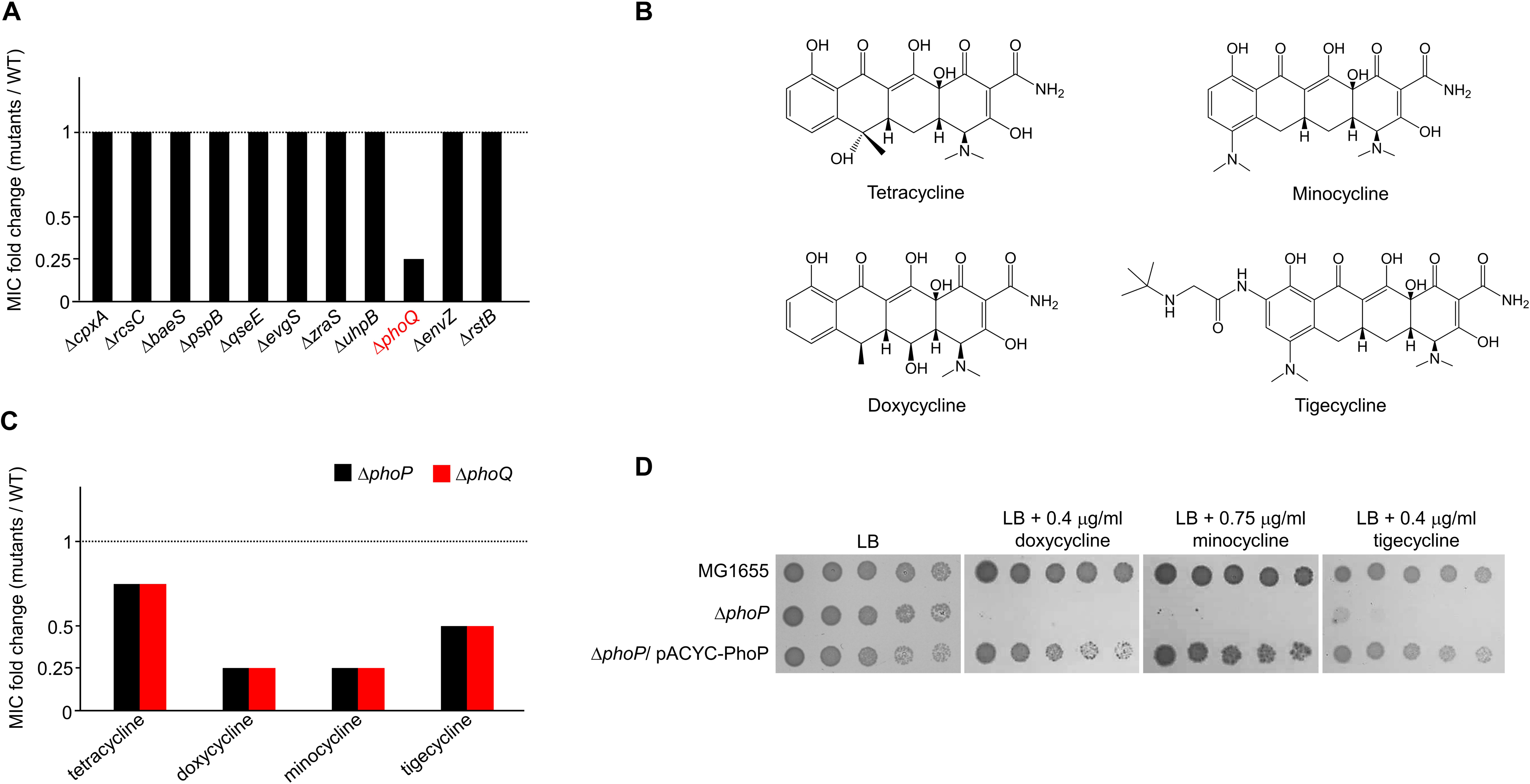
The loss of the PhoPQ two-component system confers increased susceptibility to tetracycline and glycylcycline antibiotics. (A) Increased susceptibility of the Δ*phoQ* mutant to minocycline. The MICs of minocycline were measured against the wild-type and indicated mutant strains in MH medium. The relative MIC values for the indicated mutant cells compared to those for the wild-type cells are presented. (B) Structures of tetracycline and glycylcycline antibiotics. (C) Increased susceptibility of the Δ*phoP* or Δ*phoQ* mutant to tetracycline and glycylcycline antibiotics. The MICs of indicated antibiotics were measured against the wild-type and Δ*phoP* or Δ*phoQ* mutant strains in MH medium. The relative MIC values for the Δ*phoP* (black bars) or Δ*phoQ* (red bars) mutant cells compared to those for the wild-type cells are presented. (D) Complementation of antibiotic sensitivities of the Δ*phoP* mutant. The cells of the indicated strains were serially diluted from 10^8^ to 10^4^ cells/ml in 10-fold steps and spotted onto LB plates with or without the indicated concentration of each antibiotic. The experiments were performed in triplicate, and a representative image is presented.

### PhoPQ-mediated regulation of the expression of phosphoethanolamine transferase EptB affects intrinsic resistance to doxycycline and minocycline

PhoPQ two-component system regulates the transcriptional expression of diverse genes in response to magnesium starvation and acidic or antimicrobial peptide stress (2–4, 9). The magnesium transporter MgtA is a representative member of the PhoPQ regulon, and phosphorylated PhoP activates the transcription of the *mgtA* gene (6, 7). Therefore, we examined the effect of MgtA on doxycycline and minocycline resistance. The Δ*mgtA* mutant was not sensitive to doxycycline or minocycline (Fig. 3A), and the expression of *mgtA* in the Δ*phoP* mutant did not restore its sensitivity to minocycline (Fig. 3B). These results indicate that the minocycline sensitivity of the Δ*phoP* mutant was not associated with MgtA. Phenotypic results of another magnesium transporter, CorA, were almost identical to those of MgtA (Fig. 3A and B), implying that the minocycline sensitivity of the Δ*phoP* mutant was not caused by a defect in magnesium transportation. Subsequently, doxycycline resistance effects of several additional genes (*ompT*, *borD*, *pagP*, *tolC*, and *fadL*), whose transcription was activated by phosphorylated PhoP, were tested. The expression of these genes did not restore the sensitivity of the Δ*phoP* mutant to doxycycline (Fig. 3C).

**FIG 3.**
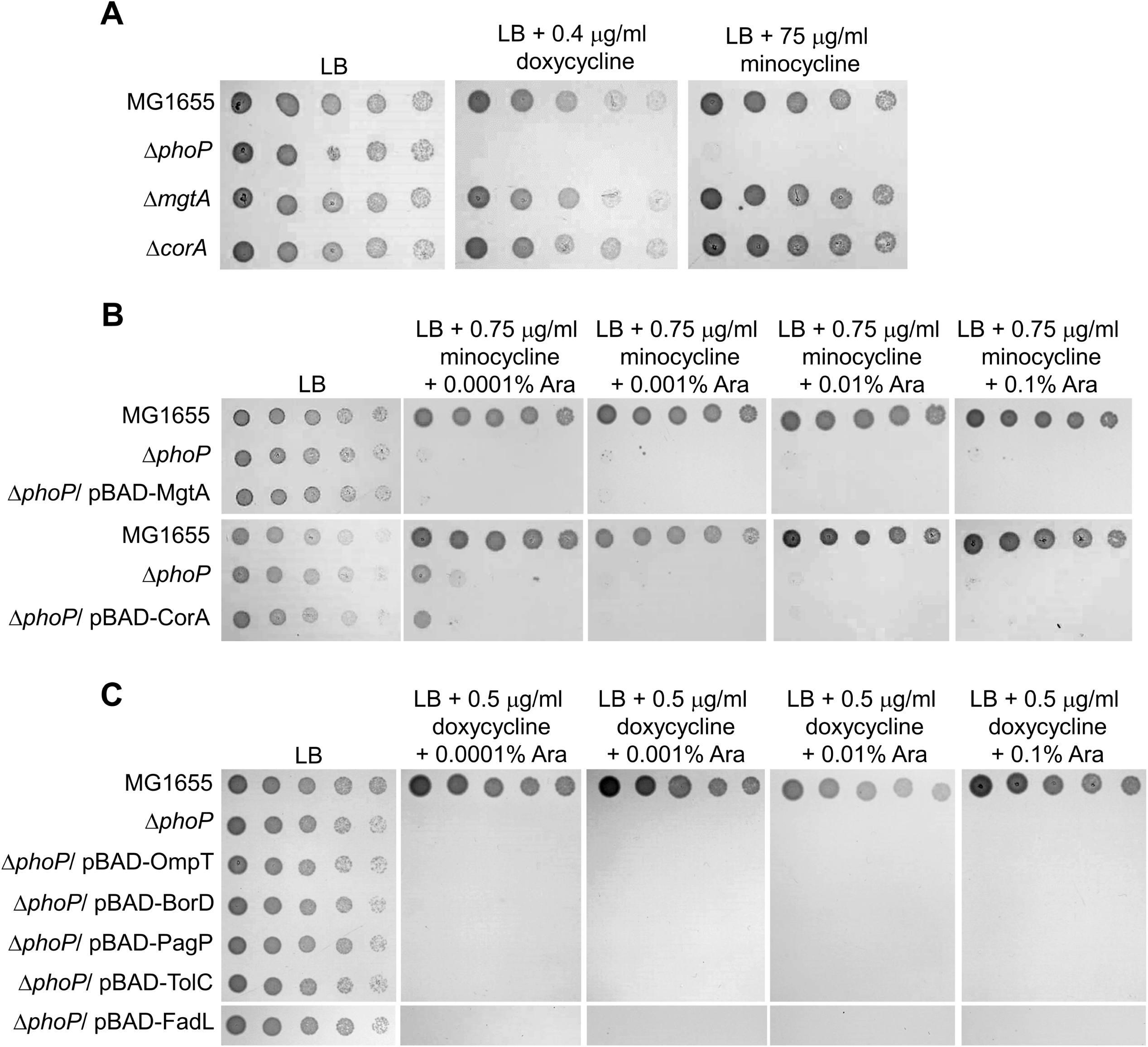
The effect of several known PhoPQ regulon genes on the resistance to tetracycline antibiotics. (A) The effect of depletion of magnesium transporter genes on doxycycline and minocycline resistance. The cells of the indicated strains were serially diluted from 10^8^ to 10^4^ cells/ml in 10-fold steps and spotted onto LB plates with or without the indicated concentration of each antibiotic. (B) The effect of expression of magnesium transporter genes on minocycline resistance in the Δ*phoP* mutant strain. The cells of the indicated strains were serially diluted from 10^8^ to 10^4^ cells/ml in 10-fold steps and spotted onto LB plates with or without the indicated concentrations of minocycline and arabinose (Ara). (C) The effect of expression of several known PhoPQ regulon genes on doxycycline resistance in the Δ*phoP* mutant strain. The cells of the indicated strains were serially diluted from 10^8^ to 10^4^ cells/ml in 10-fold steps and spotted onto LB plates with or without the indicated concentrations of doxycycline and arabinose (Ara). (A–C) The experiments were performed in triplicate, and a representative image is presented.

Finally, to identify a gene associated with doxycycline sensitivity of the Δ*phoP* mutant, we used random transposon mutagenesis to screen for suppressors in which the sensitivity of the Δ*phoP* mutant to doxycycline is restored. We isolated a suppressor in which the sensitivity to doxycycline had recovered to almost the level of the wild-type strain (Fig. 4A). The transposon insertion was mapped within *eptB* encoding the phosphoethanolamine transferase that catalyzes the addition of phosphoethanolamine to the 3-deoxy-D-manno-oct-2-ulosonate (KdoII) in the inner core of LPS (22) (Fig. 4B). We constructed a Δ*phoP* Δ*eptB* double mutant to confirm the effect of transposon insertion. The bacterial growth of this double mutant in the presence of doxycycline or minocycline was significantly restored compared to that of the Δ*phoP* mutant (Fig. 4C), thereby confirming that the deletion of the *eptB* gene suppresses the doxycycline sensitivity of the Δ*phoP* mutant. The expression of the *eptB* gene is silenced by the small regulatory RNA MgrR, whose transcription is activated by phosphorylated PhoP (23, 24). Therefore, the expression of the *eptB* gene could be activated in the Δ*phoP* mutant, which may cause doxycycline sensitivity. To assess this assumption, we measured the transcription of *eptB* and *phoP* in the wild-type and Δ*phoP* mutant strains. Expectedly, the *phoP* transcripts were not detected in the Δ*phoP* mutant, and the level of the *eptB* transcripts was almost 4-fold higher in the Δ*phoP* mutant than in the wild-type strain (Fig. 4D). Additionally, the overexpression of *eptB* in the wild-type strain using the plasmid pBAD24 induced significant sensitivity to doxycycline (Fig. 4E). Overall, these results demonstrated that PhoPQ-mediated regulation of *eptB* expression is associated with doxycycline and minocycline resistance.

**FIG 4.**
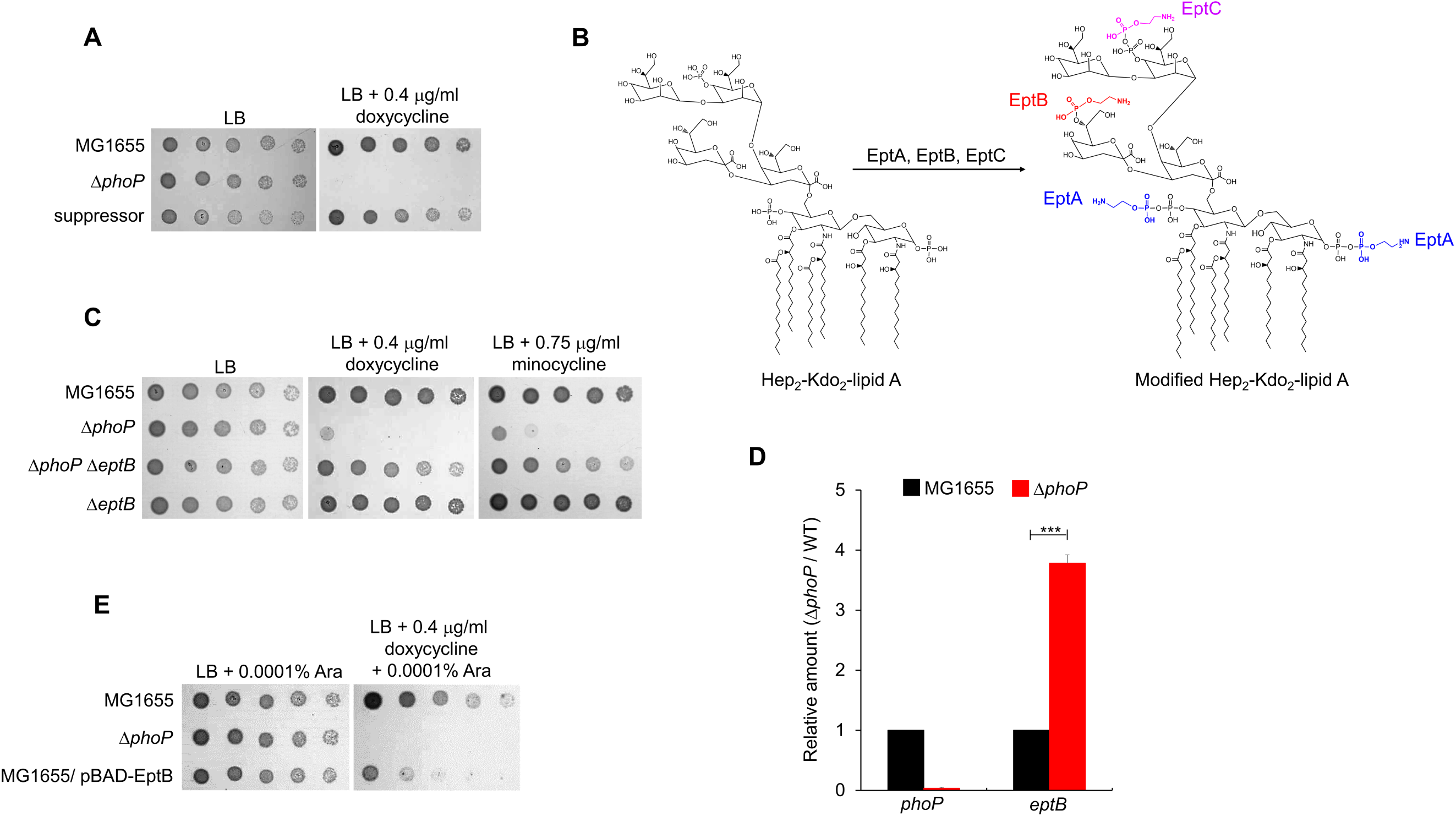
The effect of the phosphoethanolamine transferase EptB on the resistance to tetracycline antibiotics. (A) Isolation of the suppressor mutant of the Δ*phoP* mutant. The cells of the indicated strains were serially diluted from 10^8^ to 10^4^ cells/ml in 10-fold steps and spotted onto LB plates with or without the indicated concentration of doxycycline. (B) Schematic representation depicting the roles of phosphoethanolamine transferases in LPS modification. EptA, EptB, and EptC catalyze the addition of phosphoethanolamine to the phosphate group of the glucosamine disaccharide of lipid A, the KdoII sugar in the inner core, and the phosphate group of the heptose I residue in the inner core, respectively. (C) The depletion of EptB suppresses the sensitivity of the Δ*phoP* mutant to tetracycline antibiotics. The cells of the indicated strains were serially diluted from 10^8^ to 10^4^ cells/ml in 10-fold steps and spotted onto LB plates with or without the indicated concentration of doxycycline or minocycline. (D) Relative mRNA levels of the *phoP* and *eptB* genes in the wild-type and Δ*phoP* mutant strains. Total mRNA was extracted from the wild-type (black bars) and Δ*phoP* mutant (red bars) cells cultured up to the early exponential phase (OD_600_ = 0.4). mRNA levels of the *phoP* and *eptB* genes were normalized to the levels of 16S rRNA. Data were produced from three independent experiments. Statistical significance was determined using the Student’s *t*-test. \*\*\**p*<0.001. (E) The effect of overexpression of EptB on doxycycline resistance. The cells of the indicated strains were serially diluted from 10^8^ to 10^4^ cells/ml in 10-fold steps and spotted onto LB plates with or without the indicated concentrations of doxycycline and arabinose (Ara). (A, C, and E) The experiments were performed in triplicate, and a representative image is presented.

### Other phenotypes of the *phoP* mutant were not restored by the deletion of *eptB*

To assess the effect of EptB on other phenotypes of the PhoPQ two-component system, we examined additional phenotypes of the Δ*phoP* or Δ*phoQ* mutant. First, the effect of the PhoPQ two-component system on bacterial growth under various stress conditions was examined. The Δ*phoP* and Δ*phoQ* mutants were significantly sensitive to various stresses, including sodium dodecyl sulfate/EDTA, bile salt, EDTA, and acidic pH (Fig. 5A). We assessed whether the deletion of the *eptB* gene could suppress these phenotypes of the Δ*phoP* mutant, as in the case of minocycline and doxycycline. Most phenotypes of the Δ*phoP* mutant were not restored by the deletion of the *eptB* gene, although bile salt sensitivity of the Δ*phoP* mutant was slightly recovered (Fig. 5B). Next, we examined the MICs of various antibiotics against the Δ*phoP* mutant. The MICs of antibiotics for the Δ*phoP* mutant were similar to those for the Δ*phoQ* mutant (Figs. 1 and 5C). We examined the effect of EptB on antibiotic resistance in the Δ*phoP* mutant. Unlike minocycline and doxycycline, most phenotypes of the Δ*phoP* mutant against antibiotics were not restored by the deletion of the *eptB* gene, although trimethoprim sensitivity of the Δ*phoP* mutant was slightly recovered (Fig. 5D). Collectively, these results showed that most phenotypes of the Δ*phoP* mutant were not restored by the deletion of the *eptB* gene, indicating a specific relationship between EptB and the resistance to minocycline and doxycycline.

**FIG 5.**
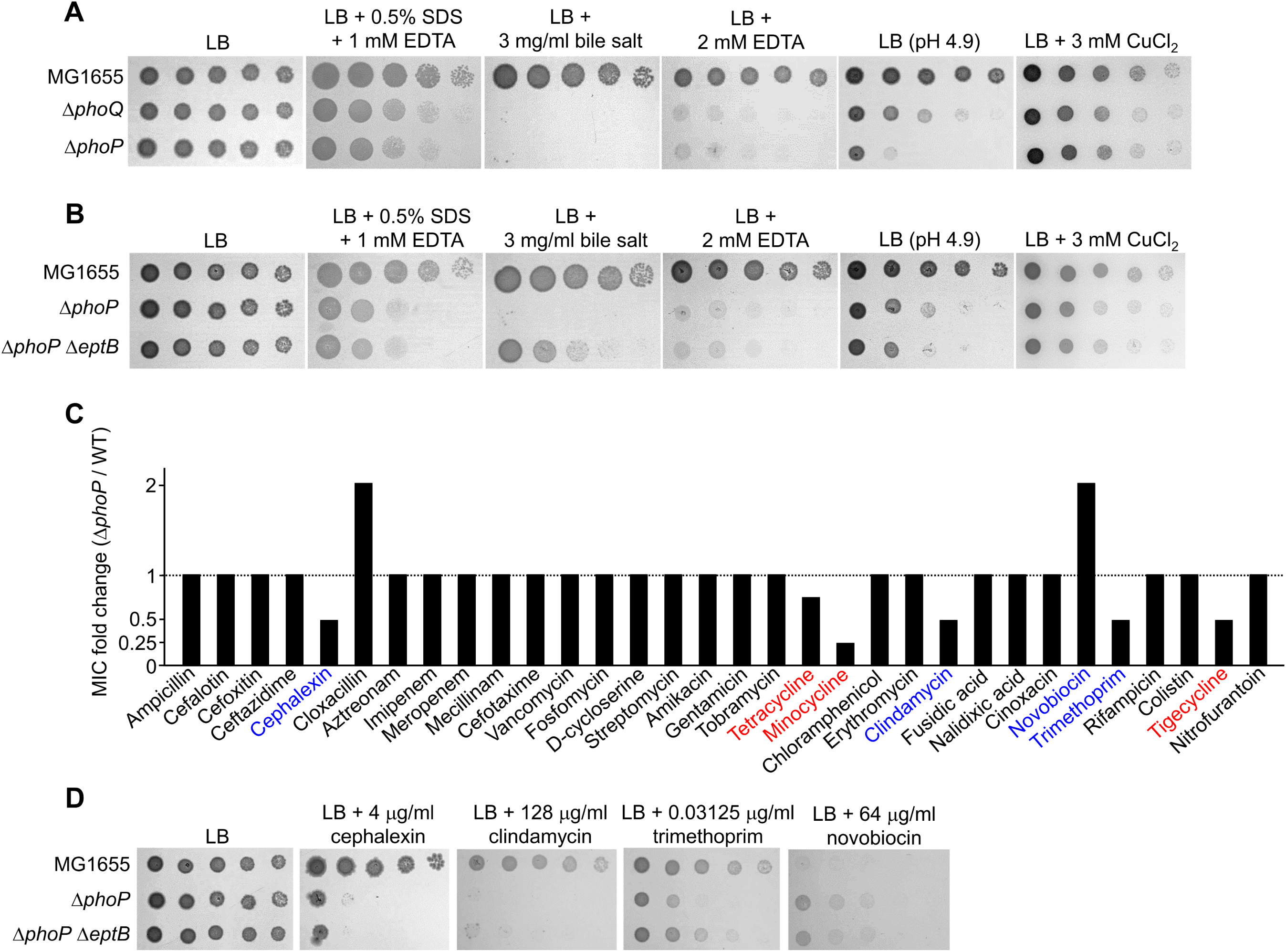
The effect of EptB on various phenotypes of the Δ*phoP* mutant. (A) Growth defect of the Δ*phoP* and Δ*phoQ* mutant strains under various envelope stress conditions. The cells of the indicated strains were serially diluted from 10^8^ to 10^4^ cells/ml in 10-fold steps and spotted onto LB plates with or without the indicated chemicals, or an acidic LB plate. (B) The effect of EptB inactivation on the sensitivity of the Δ*phoP* mutant to envelope stress. The cells of the indicated strains were serially diluted from 10^8^ to 10^4^ cells/ml in 10-fold steps and spotted onto LB plates with or without the indicated chemicals, or an acidic LB plate. (C) The effect of PhoP on the MICs of antibiotics. The MICs of various antibiotics were measured against the wild-type and Δ*phoP* mutant strains in MH medium. The relative MIC values for the Δ*phoP* mutant cells compared to those for the wild-type cells are presented. (D) The effect of EptB inactivation on the altered susceptibility of the Δ*phoP* mutant against antibiotics. The cells of the indicated strains were serially diluted from 10^8^ to 10^4^ cells/ml in 10-fold steps and spotted onto LB plates with or without the indicated concentrations of antibiotics. (A, B, and D) The experiments were performed in triplicate, and a representative image is presented.

### Depletion of EtpB alleviates increased permeability of the *phoP* mutant to doxycycline

LPS is present in the outer membrane and its modification by EptB can affect the penetration of antibiotics through the outer membrane. Therefore, we measured the penetration levels of doxycycline in the wild-type and mutant strains. Intracellular doxycycline accumulation was measured using ELISA. The Δ*phoP* mutant showed increased intracellular levels of doxycycline compared to the wild-type strain (Fig. 6). Increased levels of doxycycline in the Δ*phoP* mutant were complemented by pACYC184 plasmid-based expression of the *phoP* gene. Deletion of the *eptB* gene also diminished the increased intracellular levels of doxycycline in the Δ*phoP* mutant (Fig. 6). Therefore, these results implied that the doxycycline sensitivity of the Δ*phoP* mutant was at least partially due to the increased permeability of doxycycline.

**FIG 6.**
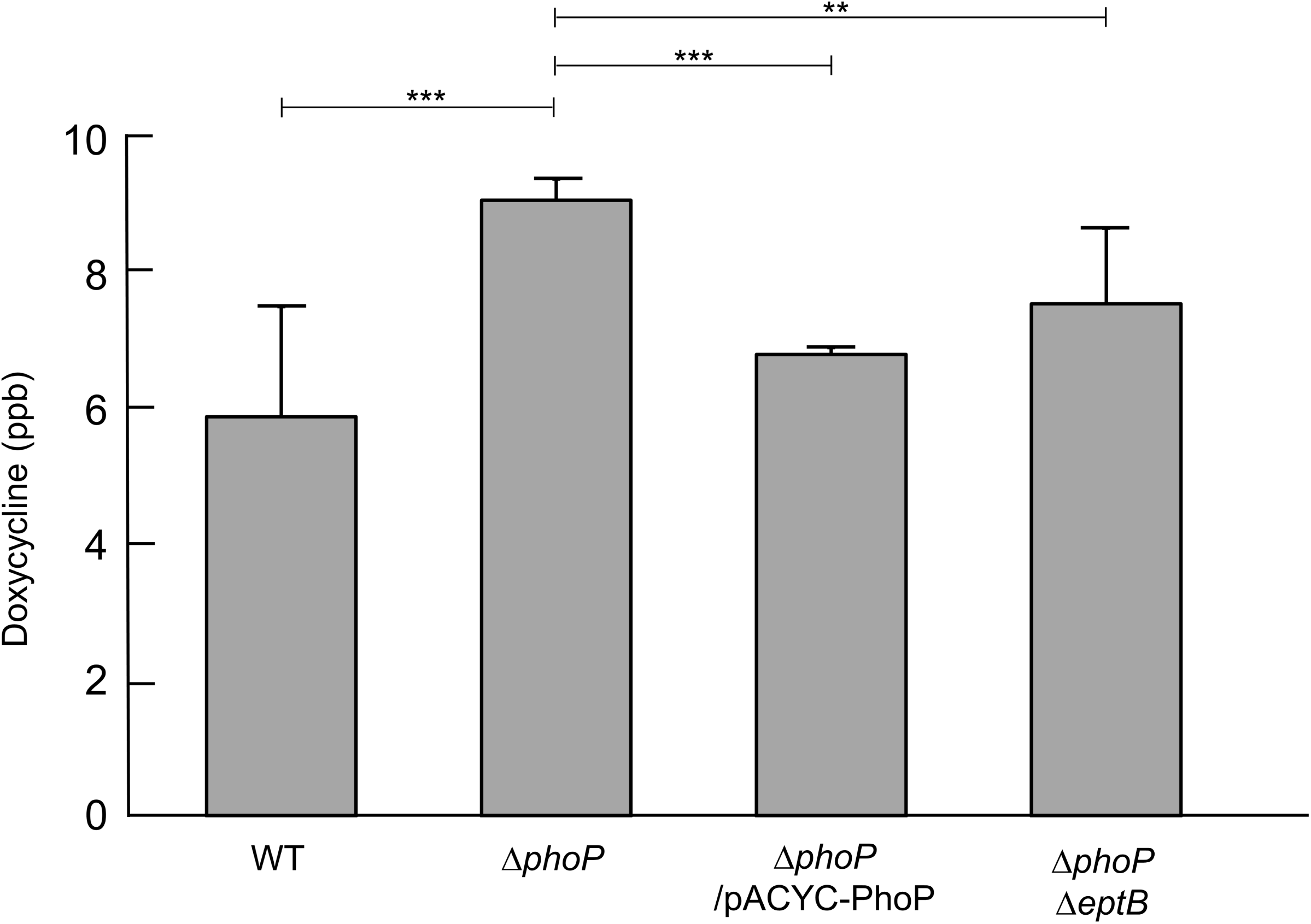
The intracellular accumulated levels of doxycycline. At the early exponential phase, 0.5 μg/ml of doxycycline was added to LB medium and the cells were harvested after additional incubation for 20 min at 37°C. After washing, the harvested cells were disrupted and cell debris was removed by centrifugation. After removing soluble proteins using acetonitrile, the doxycycline level in the supernatant was determined using a Doxycycline ELISA Kit. The doxycycline level was estimated by measuring the absorbance at 450 nm. The exact concentration of doxycycline was estimated based on the standard curve made using the standard concentrations of doxycycline. Data were produced from three independent experiments. Statistical significance was determined using the Student’s *t*-test. \*\**p*<0.01; \*\*\**p*<0.001.

### EptB is associated with resistance to tetracycline and glycylcycline antibiotics

Two other phosphoethanolamine transferases are involved in LPS modification in *E. coli*, in addition to EptB. EptA catalyzes the addition of phosphoethanolamine to the phosphate group of glucosamine disaccharide of lipid A (25, 26), whereas EptC catalyzes the incorporation of phosphoethanolamine into the phosphate group of the heptose I residue in the inner core (27) (Fig. 4B). We examined whether the deletion of *eptA* or *eptC* suppresses doxycycline sensitivity of the Δ*phoP* mutant. Unike EptB, depletion of EptA or EptC did not suppress doxycycline sensitivity of the Δ*phoP* mutant (Fig. 7A), indicating a distinct role of EptB in doxycycline resistance. We constructed single-deletion mutants of each gene and examined the MICs of various antibiotics in each mutant strain to analyze the roles of the three phosphoethanolamine transferases in more detail. Notably, neither Δ*eptA* nor Δ*eptC* mutants revealed any change in the MICs of the antibiotics tested (Fig. S3). Meanwhile, the Δ*eptB* mutant showed 2-fold and 1.5-fold higher MICs of minocycline and tigecycline, respectively, than the wild-type strain, whereas there were no changes in the MICs of other antibiotics tested (Fig. 7B). When bacterial growth of the three mutants was measured in the presence of antibiotics, the Δ*eptB* mutant exhibited resistance to minocycline, tigecycline, and doxycycline, whereas the growth levels of the Δ*eptA* and Δ*eptC* mutants were identical to those of the wild-type strain for all the antibiotics tested (Fig. 7C). In conclusion, our results demonstrate that EptB depletion induces resistance to tetracycline and glycylcycline antibiotics.

**FIG 7.**
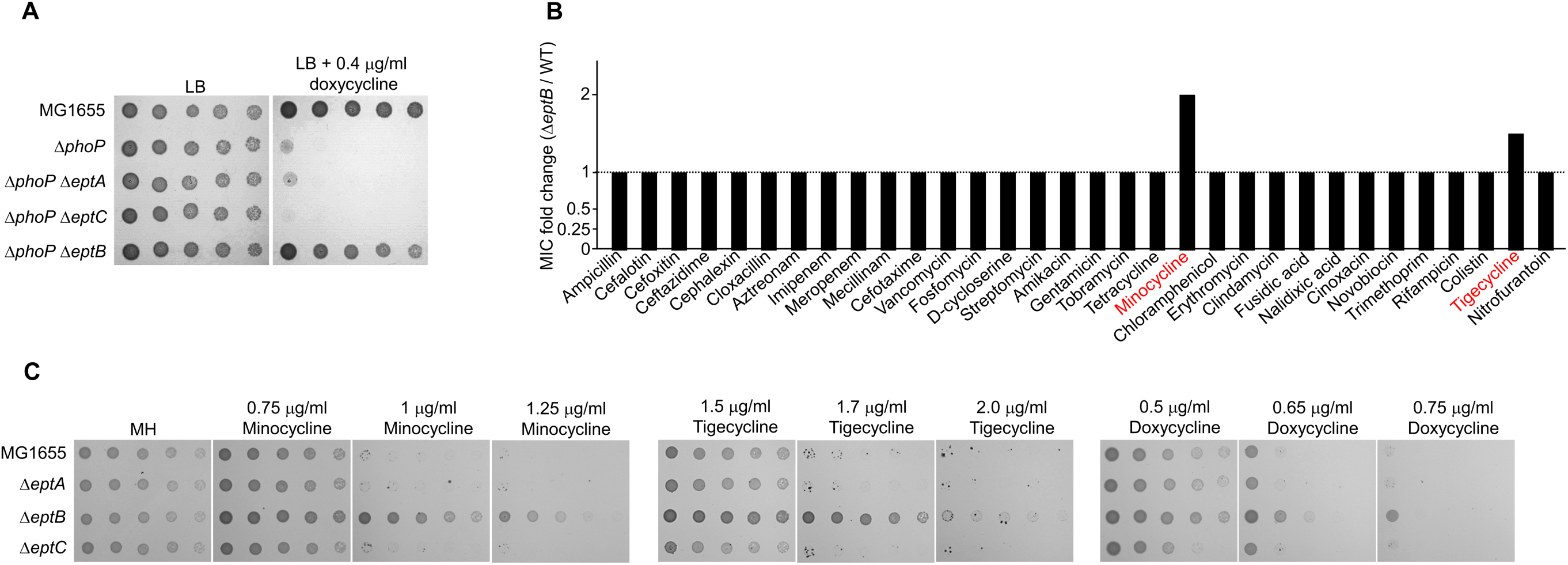
The depletion of EptB induces tetracycline and glycylcycline resistance. (A) The effect of inactivation of the phosphoethanolamine transferases on the sensitivity of the Δ*phoP* mutant to doxycycline. The cells of the indicated strains were serially diluted from 10^8^ to 10^4^ cells/ml in 10-fold steps and spotted onto LB plates with or without doxycycline. (B) The effect of EptB depletion on the MICs of antibiotics. The MICs of various antibiotics were measured against the wild-type and Δ*eptB* mutant strains in MH medium. The relative MIC values for the Δ*eptB* mutant cells compared to those for the wild-type cells are presented. (C) The resistance of the Δ*eptB* mutant to tetracycline and glycylcycline antibiotics. The cells of the indicated strains were serially diluted from 10^8^ to 10^4^ cells/ml in 10-fold steps and spotted onto LB plates with or without the indicated concentration of each antibiotic. The experiments were performed in triplicate, and a representative image is presented.

## DISCUSSION

Antibiotic resistance of Gram-negative pathogens poses a serious threat on public health worldwide (11, 28, 29). Tetracycline and glycylcycline antibiotics, especially tigecycline, are among the last resort therapeutic options for treating infections caused by multidrug-resistant Gram-negative pathogens (11, 29). Despite the clinical importance of tetracycline and glycylcycline antibiotics, the molecular mechanisms underlying the resistance to these antibiotics are not fully understood. In this study, we demonstrated that PhoPQ-mediated modification of the core region of LPS affects resistance to tetracycline and glycylcycline antibiotics. Phosphoethanolamine transferase EptB, a member of the PhoPQ regulon, adds phosphoethanolamine to the KdoII sugar in the inner core of LPS, which induces sensitivity to tetracycline and glycylcycline antibiotics (Figs. 2–4). These phenotypes appear to be caused by the increased uptake of tetracycline and glycylcycline antibiotics (Fig. 6). Modification of the inner core of LPS by EptB affected susceptibility to tetracycline and glycylcycline antibiotics, among many antibiotics with diverse modes of action (Fig. 7). These results revealed the novel physiological significance of PhoPQ-mediated modification of the inner core of LPS.

Several regions of LPS can be modified by adding a phosphoethanolamine group (22, 25–27). Among them, the addition of phosphoethanolamine to the phosphate group of the glucosamine disaccharide of lipid A has been extensively studied (30–32). Attachment of the phosphoethanolamine group neutralizes the negative charge of the lipid A phosphate group, which inhibits binding of LPS to colistin and cationic antimicrobial peptides (31–33). Consequently, this modification results in increased resistance to colistin and cationic antimicrobial peptides. The addition of other functional groups such as aminoarabinose (34), glucosamine (35), galactosamine (36, 37), and glucose (36), which can neutralize the negative charge of the lipid A phosphate group, also induces similar resistance. In contrast to lipid A modifications, studies on core modifications are scarce. Phosphoethanolamine addition to the phosphate group of heptose I in the inner core by EptC is required to overcome envelope stresses such as SDS and Zn^2+^ (27). In *E. coli,* deletion of EptC resulted in slightly increased susceptibility to polymyxin B (38). However, the physiological role of the EptB-dependent phosphoethanolamine addition to KdoII in the inner core has not yet been elucidated. Our study demonstrated that EptB overexpression increased susceptibility to tetracycline and glycylcycline antibiotics, whereas the deletion of the *eptB* gene resulted in elevated resistance to these antibiotics. These results indicate that EptB-dependent phosphoethanolamine addition to KdoII is associated with resistance to tetracycline and glycylcycline antibiotics. Notably, neither EptA nor EptC affected resistance to tetracycline and glycylcycline antibiotics, and EptB did not affect colistin resistance (Fig. 7), indicating the distinct roles among EptA, EptB, and EptC. To the best of our knowledge, this is the first report demonstrating the physiological role of KdoII modification in the inner core of LPS.

Most studies on LPS modification have been focused on the neutralization of phosphate groups present in lipid A or the core region. A reduction in the negative charge of the phosphate group of lipid A decreases the electrostatic interactions between colistin and lipid A, which induces colistin resistance (39). Meanwhile, addition of the phosphoethanolamine group to the hydroxyl group of KdoII by EptB did not induce a reduction in the formal charge of LPS (Fig. 4B). Therefore, deletion of EptB did not affect susceptibility to colistin (Fig. 7B). The MICs of all the antibiotics tested, except minocycline and tigecycline, were not changed by EptB deletion (Fig. 7B). These results indicate that EptB-mediated KdoII modification is involved in resistance to tetracycline and glycylcycline antibiotics. The mechanisms by which tetracycline and glycylcycline antibiotics penetrate the outer membrane are poorly understood. Since EptB-mediated KdoII modification affects the resistance to various tetracycline and glycylcycline antibiotics with different side-chain functional groups (Figs. 1B and 7), the penetration of a linearly fused tetracyclic nucleus across the outer membrane may be affected by KdoII modification. This aspect can be investigated in further studies.

Various resistance mechanisms against tetracycline and glycylcycline antibiotics have been unveiled, such as tetracycline-specific Tet(A) or Tet(B) efflux pump acquisition (10), GTP-dependent release mechanisms of tetracycline from the ribosome by Tet(M) or Tet(O) (10, 40, 41), and enzymatic inactivation of tetracycline by Tet(X) or Tet(37) (42, 43). Loss of the OmpF and OmpC porins induces intrinsic tetracycline resistance via reduced transportation of tetracycline across the outer membrane (44, 45). In this study, we revealed a novel intrinsic resistance mechanism for tetracycline and glycylcycline antibiotics. The phosphoethanolamine addition to the core sugar of LPS enhanced susceptibility to tetracycline and glycylcycline antibiotics (Fig. 4), whereas the loss of this modification induced intrinsic resistance to these antibiotics (Fig. 7). These effects may be caused by the altered transportation of these antibiotics across the outer membrane (Fig. 6). Therefore, our study demonstrated that LPS core modification is a novel intrinsic resistance mechanism against tetracycline and glycylcycline antibiotics.

The PhoPQ system is a two-component system that has been extensively studied in various Gram-negative bacteria (46, 47). Diverse signals that activate the sensor kinase PhoQ have been identified, including Mg^2+^ and other divalent cations (2), antimicrobial peptides (4), mildly acidic pH (3, 9), osmotic upshift (5), and long-chain unsaturated fatty acids (48). Activated response regulator PhoP regulates the transcription of many genes and diverse proteins via the regulation of other transcriptional factors, protease regulators, metabolites, and regulatory RNAs (47). These PhoP-mediated changes cause diverse phenotypic consequences in many bacterial behaviors, such as metal ion homeostasis, virulence, motility, resistance to antimicrobial agents, and the ability to overcome stressful conditions such as acidic stress or nutritional depletion (46, 47). In *E. coli*, PhoP regulates the transcription of several genes associated with LPS modification, such as *eptA* and *eptB* encoding phosphoethanolamine transferase (23, 49) and *pagP* encoding lipid A palmitoyltransferase (50). EptA induces colistin resistance via neutralization of the phosphate groups present in lipid A (31–33), whereas PagP induces resistance to cationic alpha-helical antimicrobial peptides such as C18G, magainin 2, and cecropin A (47, 51, 52). However, the physiological significance of EptB remains unclear. Here, we revealed that EptB regulates resistance to tetracycline and glycylcycline antibiotics. These results demonstrate that all PhoPQ-mediated LPS modifications are associated with antibiotic resistance.

## MATERIALS AND METHODS

### Bacterial strains, plasmids, and culture conditions

All bacterial strains and plasmids used in this study are presented in Table S1, and all primers are listed in Table S2. Bacterial cells were cultured in Luria–Bertani (LB) medium at 37°C, unless otherwise mentioned. Antibiotics, including kanamycin (50 μg/ml), chloramphenicol (5 μg/ml), tetracycline (10 μg/ml), and ampicillin (100 μg/ml), were added to the culture medium as required. Bacterial growth was estimated using a serial dilution spotting assay onto LB agar plates. Cells from overnight cultures in LB medium were inoculated into fresh LB medium. On reaching an OD_600_ of approximately 0.8, the cultured cells were serially diluted 10-fold from 10^8^ to 10^4^ cells/ml, and 2 μl of the samples were spotted onto LB agar plates with or without the indicated chemicals. After incubation at 37°C for 10–20 h, plates were imaged using a digital camera, EOS 100D (Canon Inc., Japan).

All deletion mutants were constructed using λ red recombinase, as described previously (53). DNA products for gene deletion were prepared by polymerase chain reaction (PCR) using primers with 50 bp sequence for homologous recombination, and the plasmid pKD13 with a kanamycin resistance gene as a template. After purification of the PCR products, the purified deletion cassettes were electroporated into MG1655 or mutant cells harboring the plasmid pKD46 expressing λ red recombinase. Deletion mutants were selected on LB plates containing kanamycin, and the deletion of the target gene was confirmed using PCR. The kanamycin resistance gene inserted into the chromosome was removed by the plasmid pCP20 expressing the FLP recombinase, as described previously (53). The plasmid pCP20 in the mutant cells was removed by incubation at 37°C, instead of incubation at 42°C, to decrease physiological changes in the bacteria cells (54, 55).

DNA covering both the promoter region and open reading frame of PhoP (from −170 to +690) was cloned into the plasmid pACYC184. PCR was performed by using a forward primer possessing a synthetic BamHI site (underlined) (5′-CCCGTCCTGT<u>GGATCC</u>AAACCTCGTATCAGTGCCGG-3′), and a reverse primer possessing a synthetic EagI site (underlined) (5′-CCCAGCGCGT<u>CGGCCG</u>GACGCAGTAATTTTTTCATC-3′). PCR product was inserted into the plasmid pACYC184 digested using BamHI and EagI by infusion cloning (Clontech, USA), as reported previously (56). Cloning was confirmed by PCR using other primer sets located within the plasmid pACYC184, and DNA sequencing. A pBAD24 plasmid expressing genes regulated by the PhoPQ system, such as *mgtA*, was constructed using a similar method. The entire open reading frame of each gene was amplified using PCR, and the PCR product was inserted into the plasmid pBAD24 digested by EcoRI and XbaI, through homologous recombination between overlapping sequences using infusion cloning. Target gene cloning was confirmed using DNA sequencing.

### Determination of minimal inhibitory concentrations (MICs) of antibiotics

The MICs of antibiotics were determined according to the guidelines provided by the Clinical and Laboratory Standards Institute (57). All wild-type and mutant strains were cultured overnight in Mueller–Hinton (MH) broth and subsequently inoculated into fresh MH medium. When the bacterial suspensions reached a turbidity of 0.5 McFarland standard (approximately 1.5 × 10^8^ cells/ml), the cells were diluted to a final concentration of 10^7^ cells/ml using MH broth. A diluted suspension of 10 μl was spotted onto MH plates containing antibiotics at final concentrations ranging from 1024 μg/ml to 7.8 ng/ml in two-fold serial dilutions. The MIC of each antibiotic was determined after incubation at 37°C for 20 h, based on the bacterial growth. The MIC corresponds to the lowest concentration at which visible lawn growth of the cell spot is inhibited.

### Transposon mutagenesis and identification of transposon insertion site

Transposon mutagenesis was performed to identify a mutant that suppresses the doxycycline sensitivity of the Δ*phoP* mutant, using the *pir*-dependent transposon delivery vector pRL27 carrying a Tn5 transposase gene and a mini-Tn5 element encoding kanamycin resistance (58). The pRL27 plasmid was amplified in *E. coli* DH5αλ*pir* cells carrying the *pir* gene. Purified pRL27 plasmids were electroporated into competent cells of the Δ*phoP* mutant. The pRL27 plasmid was not replicated in this strain as the Δ*phoP* mutant did not harbor the *pir* gene; therefore, the kanamycin resistance gene in this strain was maintained when the chromosomal insertion of mini-Tn5 element occurred. The mutant that suppressed doxycycline sensitivity of the Δ*phoP* mutant was selected using an LB plate containing both kanamycin (50 μg/ml) and doxycycline (0.4 μg/ml). PCR was performed to uncover the transposon insertion site, using the genomic DNA of the suppressor strain as a template and primer sequences (an arbitrary primer consisting of a GGCGGT sequence and a random sequence, and a Tn5 transposon inner primer, 5′-GGTTGTAACACTGGCAGAGCATTACG-3′), as described previously (59). After PCR purification, product was sequenced using another Tn5 transposon inner primer, 5′-ATCAGCAACTTAAATAGCCTCTAAGG-3′.

### Quantitative real-time RT-PCR

Wild-type and Δ*phoP* mutant strains cultured overnight in LB medium were inoculated into fresh LB medium and cultured at 37°C to reach early exponential phase. Total RNA was extracted from the cells using the RNeasy Mini Kit (Qiagen, USA). Contaminating DNA in the samples were removed through incubation at 37°C for 2 h using RNase-free DNase I (Promega, USA). All RNAs in the samples were converted into cDNA using a cDNA EcoDry Premix (Clontech, USA). cDNA levels of the *phoP* and *eptB* genes were quantified by PCR using SYBR Premix Ex Taq II (Takara, Japan) solution containing RT-PCR primers for each gene (See Table S2) and 10-fold diluted cDNA samples as templates, in a CFX96 Real-Time System (Bio-Rad, USA). The 16S rRNA gene was used as the reference to estimate the expression level of each gene.

### Estimation of doxycycline uptake

Cells cultured overnight in LB medium were inoculated into fresh LB medium. At the early exponential phase, 0.5 μg/ml of doxycycline was added into the LB medium and cells were cultured for 20 min at 37°C. Cells were harvested and washed using washing buffer (100 mM Tris-HCl [pH 8.0], and 100 mM NaCl). The cells were resuspended in 1 ml of resuspension buffer (50 mM Tris-HCl [pH 8.0], and 300 mM NaCl) and disrupted using a French press at 8,000 psi. The sample was centrifuged at 10,000 × *g* for 10 min at 4°C, and the supernatant was mixed with 1 ml of acetonitrile. Precipitated proteins were removed by centrifugation at 10,000 × *g* for 5 min at 4°C and the supernatant was diluted 5-fold using distilled water. Doxycycline levels in diluted samples were determined using a Doxycycline ELISA Kit (BioVision, USA). Doxycycline levels were estimated by measuring the absorbance at 450 nm. The exact concentration of doxycycline was estimated based on a standard curve of doxycycline drawn using standard concentrations.

## ACKNOWLEDGMENTS

This work was supported by research grants from Basic Science Research Program through the National Research Foundation of Korea funded by the Ministry of Education (NRF-RS-2023-00246684 and 2021R1A6A3A01086629) and Korea Institute of Planning and Evaluation for Technology in Food, Agriculture and Forestry through High Value-added Food Technology Development Program, funded by Ministry of Agriculture, Food and Rural Affairs (grant number 322026-3).

## REFERENCES

1. Zschiedrich CP, Keidel V, Szurmant H. 2016. Molecular mechanisms of two-component signal transduction. J Mol Biol 428:3752–3775. 10.1016/j.jmb.2016.08.003.

2. Garcia Vescovi E, Soncini FC, Groisman EA. 1996. Mg^2+^ as an extracellular signal: environmental regulation of *Salmonella* virulence. Cell 84:165–174. 10.1016/s0092-8674(00)81003-x.

3. Prost LR, Daley ME, Le Sage V, Bader MW, Le Moual H, Klevit RE, Miller SI. 2007. Activation of the bacterial sensor kinase PhoQ by acidic pH. Mol Cell 26:165–174. 10.1016/j.molcel.2007.03.008.

4. Bader MW, Sanowar S, Daley ME, Schneider AR, Cho U, Xu W, Klevit RE, Le Moual H, Miller SI. 2005. Recognition of antimicrobial peptides by a bacterial sensor kinase. Cell 122:461–472. 10.1016/j.cell.2005.05.030.

5. Yuan J, Jin F, Glatter T, Sourjik V. 2017. Osmosensing by the bacterial PhoQ/PhoP two-component system. Proc Natl Acad Sci U S A 114:E10792–E10798. 10.1073/pnas.1717272114.

6. Minagawa S, Ogasawara H, Kato A, Yamamoto K, Eguchi Y, Oshima T, Mori H, Ishihama A, Utsumi R. 2003. Identification and molecular characterization of the Mg^2+^ stimulon of *Escherichia coli*. J Bacteriol 185:3696–3702. 10.1128/JB.185.13.3696-3702.2003.

7. Yamamoto K, Ogasawara H, Fujita N, Utsumi R, Ishihama A. 2002. Novel mode of transcription regulation of divergently overlapping promoters by PhoP, the regulator of two-component system sensing external magnesium availability. Mol Microbiol 45:423–438. 10.1046/j.1365-2958.2002.03017.x.

8. Lee EJ, Pontes MH, Groisman EA. 2013. A bacterial virulence protein promotes pathogenicity by inhibiting the bacterium’s own F1Fo ATP synthase. Cell 154:146–156. 10.1016/j.cell.2013.06.004.

9. Alpuche Aranda CM, Swanson JA, Loomis WP, Miller SI. 1992. *Salmonella typhimurium* activates virulence gene transcription within acidified macrophage phagosomes. Proc Natl Acad Sci U S A 89:10079–10083. 10.1073/pnas.89.21.10079.

10. Grossman TH. 2016. Tetracycline antibiotics and resistance. Cold Spring Harb Perspect Med 6:a025387. 10.1101/cshperspect.a025387.

11. Yaghoubi S, Zekiy AO, Krutova M, Gholami M, Kouhsari E, Sholeh M, Ghafouri Z, Maleki F. 2022. Tigecycline antibacterial activity, clinical effectiveness, and mechanisms and epidemiology of resistance: narrative review. Eur J Clin Microbiol Infect Dis 41:1003–1022. 10.1007/s10096-020-04121-1.

12. Pankey GA. 2005. Tigecycline. J Antimicrob Chemother 56:470–480. 10.1093/jac/dki248.

13. Sun C, Yu Y, Hua X. 2023. Resistance mechanisms of tigecycline in *Acinetobacter baumannii*. Front Cell Infect Microbiol 13:1141490. 10.3389/fcimb.2023.1141490.

14. Zhang S, Wen J, Wang Y, Wang M, Jia R, Chen S, Liu M, Zhu D, Zhao X, Wu Y, Yang Q, Huang J, Ou X, Mao S, Gao Q, Sun D, Tian B, Cheng A. 2022. Dissemination and prevalence of plasmid-mediated high-level tigecycline resistance gene tet (X4). Front Microbiol 13:969769. 10.3389/fmicb.2022.969769.

15. Speer BS, Shoemaker NB, Salyers AA. 1992. Bacterial resistance to tetracycline: mechanisms, transfer, and clinical significance. Clin Microbiol Rev 5:387–399. 10.1128/CMR.5.4.387.

16. Masi M, Pinet E, Pages JM. 2020. Complex response of the CpxAR two-component system to β-lactams on antibiotic resistance and envelope homeostasis in *Enterobacteriaceae*. Antimicrob Agents Chemother 64 10.1128/AAC.00291-20.

17. Choi U, Lee CR. 2019. Distinct roles of outer membrane porins in antibiotic resistance and membrane integrity in *Escherichia coli*. Front Microbiol 10:953. 10.3389/fmicb.2019.00953.

18. Castaneda-Garcia A, Blazquez J, Rodriguez-Rojas A. 2013. Molecular mechanisms and clinical impact of acquired and intrinsic fosfomycin resistance. Antibiotics (Basel) 2:217–236. 10.3390/antibiotics2020217.

19. Munro LJ, Kell DB. 2022. Analysis of a library of *Escherichia coli* transporter knockout strains to identify transport pathways of antibiotics. Antibiotics (Basel) 11 10.3390/antibiotics11081129.

20. Rafailidis P, Panagopoulos P, Koutserimpas C, Samonis G. 2024. Current therapeutic approaches for multidrug-resistant and extensively drug-resistant *Acinetobacter baumannii* infections. Antibiotics (Basel) 13 10.3390/antibiotics13030261.

21. Bishburg E, Bishburg K. 2009. Minocycline-an old drug for a new century: emphasis on methicillin-resistant *Staphylococcus aureus* (MRSA) and *Acinetobacter baumannii*. Int J Antimicrob Agents 34:395–401. 10.1016/j.ijantimicag.2009.06.021.

22. Reynolds CM, Kalb SR, Cotter RJ, Raetz CR. 2005. A phosphoethanolamine transferase specific for the outer 3-deoxy-D-manno-octulosonic acid residue of *Escherichia coli* lipopolysaccharide. Identification of the *eptB* gene and Ca^2+^ hypersensitivity of an *eptB* deletion mutant. J Biol Chem 280:21202–21211. 10.1074/jbc.M500964200.

23. Moon K, Gottesman S. 2009. A PhoQ/P-regulated small RNA regulates sensitivity of *Escherichia coli* to antimicrobial peptides. Mol Microbiol 74:1314–1330. 10.1111/j.1365-2958.2009.06944.x.

24. Moon K, Six DA, Lee HJ, Raetz CR, Gottesman S. 2013. Complex transcriptional and post-transcriptional regulation of an enzyme for lipopolysaccharide modification. Mol Microbiol 89:52–64. 10.1111/mmi.12257.

25. Purcell AB, Voss BJ, Trent MS. 2022. Diacylglycerol kinase A is essential for polymyxin resistance provided by EptA, MCR-1, and other lipid A phosphoethanolamine transferases. J Bacteriol 204:e0049821. 10.1128/JB.00498-21.

26. Herrera CM, Hankins JV, Trent MS. 2010. Activation of PmrA inhibits LpxT-dependent phosphorylation of lipid A promoting resistance to antimicrobial peptides. Mol Microbiol 76:1444–1460. 10.1111/j.1365-2958.2010.07150.x.

27. Klein G, Muller-Loennies S, Lindner B, Kobylak N, Brade H, Raina S. 2013. Molecular and structural basis of inner core lipopolysaccharide alterations in *Escherichia coli*: incorporation of glucuronic acid and phosphoethanolamine in the heptose region. J Biol Chem 288:8111–8127. 10.1074/jbc.M112.445981.

28. Lee CR, Cho IH, Jeong BC, Lee SH. 2013. Strategies to minimize antibiotic resistance. Int J Environ Res Public Health 10:4274–4305. 10.3390/ijerph10094274.

29. Lee CR, Lee JH, Park M, Park KS, Bae IK, Kim YB, Cha CJ, Jeong BC, Lee SH. 2017. Biology of *Acinetobacter baumannii*: pathogenesis, antibiotic resistance mechanisms, and prospective treatment options. Front Cell Infect Microbiol 7:55. 10.3389/fcimb.2017.00055.

30. Xu Y, Wei W, Lei S, Lin J, Srinivas S, Feng Y. 2018. An evolutionarily conserved mechanism for intrinsic and transferable polymyxin resistance. mBio 9 10.1128/mBio.02317-17.

31. Raetz CR, Reynolds CM, Trent MS, Bishop RE. 2007. Lipid A modification systems in gram-negative bacteria. Annu Rev Biochem 76:295–329. 10.1146/annurev.biochem.76.010307.145803.

32. Simpson BW, Trent MS. 2019. Pushing the envelope: LPS modifications and their consequences. Nat Rev Microbiol 17:403–416. 10.1038/s41579-019-0201-x.

33. Nummila K, Kilpelainen I, Zahringer U, Vaara M, Helander IM. 1995. Lipopolysaccharides of polymyxin B-resistant mutants of *Escherichia coli* are extensively substituted by 2-aminoethyl pyrophosphate and contain aminoarabinose in lipid A. Mol Microbiol 16:271–278. 10.1111/j.1365-2958.1995.tb02299.x.

34. Trent MS, Ribeiro AA, Lin S, Cotter RJ, Raetz CR. 2001. An inner membrane enzyme in *Salmonella* and *Escherichia coli* that transfers 4-amino-4-deoxy-L-arabinose to lipid A: induction on polymyxin-resistant mutants and role of a novel lipid-linked donor. J Biol Chem 276:43122–43131. 10.1074/jbc.M106961200.

35. Rolin O, Muse SJ, Safi C, Elahi S, Gerdts V, Hittle LE, Ernst RK, Harvill ET, Preston A. 2014. Enzymatic modification of lipid A by ArnT protects *Bordetella bronchiseptica* against cationic peptides and is required for transmission. Infect Immun 82:491–499. 10.1128/IAI.01260-12.

36. Song F, Guan Z, Raetz CR. 2009. Biosynthesis of undecaprenyl phosphate-galactosamine and undecaprenyl phosphate-glucose in *Francisella novicida*. Biochemistry 48:1173–1182. 10.1021/bi802212t.

37. Wang X, Ribeiro AA, Guan Z, Raetz CR. 2009. Identification of undecaprenyl phosphate-β-D-galactosamine in *Francisella novicida* and its function in lipid A modification. Biochemistry 48:1162–1172. 10.1021/bi802211k.

38. Salazar J, Alarcon M, Huerta J, Navarro B, Aguayo D. 2017. Phosphoethanolamine addition to the heptose I of the lipopolysaccharide modifies the inner core structure and has an impact on the binding of polymyxin B to the *Escherichia coli* outer membrane. Arch Biochem Biophys 620:28–34. 10.1016/j.abb.2017.03.008.

39. Jeannot K, Bolard A, Plesiat P. 2017. Resistance to polymyxins in Gram-negative organisms. Int J Antimicrob Agents 49:526–535. 10.1016/j.ijantimicag.2016.11.029.

40. Connell SR, Tracz DM, Nierhaus KH, Taylor DE. 2003. Ribosomal protection proteins and their mechanism of tetracycline resistance. Antimicrob Agents Chemother 47:3675–3681. 10.1128/AAC.47.12.3675-3681.2003.

41. Connell SR, Trieber CA, Dinos GP, Einfeldt E, Taylor DE, Nierhaus KH. 2003. Mechanism of Tet(O)-mediated tetracycline resistance. EMBO J 22:945–953. 10.1093/emboj/cdg093.

42. Yang W, Moore IF, Koteva KP, Bareich DC, Hughes DW, Wright GD. 2004. TetX is a flavin-dependent monooxygenase conferring resistance to tetracycline antibiotics. J Biol Chem 279:52346–52352. 10.1074/jbc.M409573200.

43. Diaz-Torres ML, McNab R, Spratt DA, Villedieu A, Hunt N, Wilson M, Mullany P. 2003. Novel tetracycline resistance determinant from the oral metagenome. Antimicrob Agents Chemother 47:1430–1432. 10.1128/AAC.47.4.1430-1432.2003.

44. Mortimer PG, Piddock LJ. 1993. The accumulation of five antibacterial agents in porin-deficient mutants of *Escherichia coli*. J Antimicrob Chemother 32:195–213. 10.1093/jac/32.2.195.

45. Thanassi DG, Suh GS, Nikaido H. 1995. Role of outer membrane barrier in efflux-mediated tetracycline resistance of *Escherichia coli*. J Bacteriol 177:998–1007. 10.1128/jb.177.4.998-1007.1995.

46. Groisman EA. 2001. The pleiotropic two-component regulatory system PhoP-PhoQ. J Bacteriol 183:1835–1842. 10.1128/JB.183.6.1835-1842.2001.

47. Groisman EA, Duprey A, Choi J. 2021. How the PhoP/PhoQ system controls virulence and Mg^2+^ homeostasis: lessons in signal transduction, pathogenesis, physiology, and evolution. Microbiol Mol Biol Rev 85:e0017620. 10.1128/MMBR.00176-20.

48. Viarengo G, Sciara MI, Salazar MO, Kieffer PM, Furlan RL, Garcia Vescovi E. 2013. Unsaturated long chain free fatty acids are input signals of the *Salmonella enterica* PhoP/PhoQ regulatory system. J Biol Chem 288:22346–22358. 10.1074/jbc.M113.472829.

49. Rubin EJ, Herrera CM, Crofts AA, Trent MS. 2015. PmrD is required for modifications to *Escherichia coli* endotoxin that promote antimicrobial resistance. Antimicrob Agents Chemother 59:2051–2061. 10.1128/AAC.05052-14.

50. Eguchi Y, Okada T, Minagawa S, Oshima T, Mori H, Yamamoto K, Ishihama A, Utsumi R. 2004. Signal transduction cascade between EvgA/EvgS and PhoP/PhoQ two-component systems of *Escherichia coli*. J Bacteriol 186:3006–3014. 10.1128/JB.186.10.3006-3014.2004.

51. Guo L, Lim KB, Poduje CM, Daniel M, Gunn JS, Hackett M, Miller SI. 1998. Lipid A acylation and bacterial resistance against vertebrate antimicrobial peptides. Cell 95:189–198. 10.1016/s0092-8674(00)81750-x.

52. Shi Y, Cromie MJ, Hsu FF, Turk J, Groisman EA. 2004. PhoP-regulated *Salmonella* resistance to the antimicrobial peptides magainin 2 and polymyxin B. Mol Microbiol 53:229–241. 10.1111/j.1365-2958.2004.04107.x.

53. Datsenko KA, Wanner BL. 2000. One-step inactivation of chromosomal genes in *Escherichia coli* K-12 using PCR products. Proc Natl Acad Sci U S A 97:6640–6645. 10.1073/pnas.120163297.

54. Park SH, Kim YJ, Lee HB, Seok YJ, Lee CR. 2020. Genetic evidence for distinct functions of peptidoglycan endopeptidases in *Escherichia coli*. Front Microbiol 11:565767. 10.3389/fmicb.2020.565767.

55. Son JE, Park SH, Choi U, Lee CR. 2024. Lytic transglycosylase repertoire diversity enables intrinsic antibiotic resistance and daughter cell separation in *Escherichia coli* under acidic stress. Antimicrob Agents Chemother:e0037224. 10.1128/aac.00372-24.

56. Choi U, Park SH, Lee HB, Son JE, Lee CR. 2023. Coordinated and distinct roles of peptidoglycan carboxypeptidases DacC and DacA in cell growth and shape maintenance under stress conditions. Microbiol Spectr 11:e0001423. 10.1128/spectrum.00014-23.

57. Wikler MA, CLSI. 2018. Methods for dilution antimicrobial susceptibility tests for bacteria that grow aerobically; approved standard. 11th edn Wayne, PA: Clinical and Laboratory Standards Institute.

58. Larsen RA, Wilson MM, Guss AM, Metcalf WW. 2002. Genetic analysis of pigment biosynthesis in *Xanthobacter autotrophicus* Py2 using a new, highly efficient transposon mutagenesis system that is functional in a wide variety of bacteria. Arch Microbiol 178:193–201. 10.1007/s00203-002-0442-2.

59. Lee HB, Park SH, Lee CR. 2021. The inner membrane protein LapB is required for adaptation to cold stress in an LpxC-independent manner. J Microbiol 59:666–674. 10.1007/s12275-021-1130-8.

